# Primate mothers promote proximity between their offspring and infants who look like them

**DOI:** 10.1101/2022.05.09.491128

**Authors:** Marie J. E. Charpentier, Clémence Poirotte, Berta Roura-Torres, Paul Amblard-Rambert, Eric Willaume, Peter M. Kappeler, François Rousset, Julien P. Renoult

## Abstract

Behavioral discrimination of kin is a key process structuring social relationships in animals. In this study, we provide a first example of discrimination towards non-kin by third-parties through a mechanism of phenotype matching. In mandrills, we recently demonstrated increased facial resemblance among paternally-related juvenile and adult females indicating adaptive opportunities for paternal kin recognition. Here, we hypothesize that mothers use offspring’s facial resemblance with other infants to guide offspring’s social opportunities towards similarly-looking ones. Using deep learning for face recognition in 80 wild mandrill infants, we first show that infants born to the same father or conceived during the tenure of the same alpha male resemble each other the most, independently of their age, sex or maternal origin, extending previous results to the youngest age class. Using long-term behavioral observations on association patterns and controlling for matrilineal origin, maternal relatedness and infant age and sex, we then demonstrate that, as hypothesized, mothers are spatially closer to infants that resemble their own offspring more, thereby facilitating associations among similar-looking infants. Using theoretical modeling, we describe a plausible evolutionary process whereby mothers gain fitness benefits by promoting nepotism among paternally related infants. This mechanism, that we call “second-order kin selection”, may extend beyond mother-infant interactions and has the potential to explain cooperative behaviors among non-kin in social species, including humans.

## Introduction

Kin selection is an evolutionary process promoting traits that provide fitness benefits for genetic relatives of individuals expressing them^1^. Empirical observations of diverse interactions arising from kin selection have been pervasively reported *in natura* and constitute the foundations of many studies on social evolution^2^. Kin selection often requires kin recognition, which can operate through phenotype matching^3^. This mechanism, based on learning processes of odors, sounds or visual cues, allows individuals to recognize kin based on phenotypic resemblance either with self (“self-referent phenotype matching”^4,5^) or with other kin used as templates^6^. Self-referent phenotype matching requires individuals to evaluate their own phenotype (e.g. through smell in rodents^5,7^), which may be challenging in some situations. For example, although face recognition is a crucial prerequisite for visual communication and therefore for the maintenance of social relationships in many species^8^, including in our lineage (e.g.^9–12^), in natural contexts, animals other than humans have probably limited access to cues regarding their own facial traits (but see^4,13^). Using familiar kin as templates to recognize unfamiliar kin also requires particular conditions, including the presence of relatives during template formation. This mechanism has been rarely demonstrated in the wild (but see^14^ and^15,16^ for lab studies). A crucial question is therefore how an individual may recognize unfamiliar kin when it cannot match phenotypes to itself or to other templates.

In mandrills (*Mandrillus sphinx*), an Old World primate from Central Africa, we have recently suggested that kin recognition by phenotype matching could emerge from third-parties promoting interactions between individuals with similar faces^17^. Mandrills live in large matrilineal societies characterized by family units of philopatric, maternally-related and highly nepotistic females. Males, the dispersing sex, are only temporary residents in these social groups and the highest-ranking male sires a large proportion of infants born into different matrilines each year^17,18^. Each cohort of new-borns therefore includes a large proportion of paternal half-sibs (PHS). Nepotism is observed among these PHS, as early as juvenility^19^ and until adulthood, at least among the philopatric female mandrills^17^. Previously, we have demonstrated that unfamiliar kin recognize each other using phenotype matching based on acoustic^14^ and possibly visual cues^20^, and that facial resemblance correlates positively with genetic relatedness across female mandrills^17^. Crucially, female PHS resemble each other visually more than maternal half-sisters (MHS) do^17^, even though both kin categories share, on average, similar degrees of genetic relatedness (r~0.25; and see Supplementary Information S1). This heightened facial resemblance among PHS compared to MHS, possibly resulting from genomic imprinting processes^21,22^, indicates adaptive opportunities for paternal kin recognition, necessarily mediated by phenotype matching mechanisms. Using self-referent phenotype matching to discriminate PHS appears, however, unrealistic in wild mandrills because of environmental constraints (no physical medium to allow facial self-recognition). Face similarity could also result from selection processes on other self-evaluable phenotypic traits such as body odors, but this mechanism fails to explain the *increased* facial resemblance observed among PHS (Supplementary Information S2a). Using the father’s face as a template is also unlikely in mandrills because of the highly pronounced sexual dimorphism and morphological differences between adults and immatures in this species (infants do not resemble their father: Supplementary Information S3 and Table S1a). In Charpentier et al. (2020), we discussed an alternative mechanism where third-parties would use increased facial resemblance among PHS to shape their social relationships. We proposed that mothers could evaluate their offspring’s facial resemblance with other youngsters and guide their offspring’s social opportunities towards similar-looking ones, paving the way for nepotism among PHS. If mothers indeed manipulate their offspring’s social preferences, we expect this to occur very early during development, in infants aged ≤ 1 yr, for two reasons. First, in nonhuman primates, the first year of life represents a developmental stage where infants are still under strong maternal control^23^. Second, differences in facial resemblance among PHS compared to MHS or non-kin were found higher during juvenescence (2-4 yrs) than during adulthood^17^ in this species.

In this study, we provide the first empirical evidence that mothers drive spatial association among similarly-looking infants, who are more likely than expected by chance to be paternally-related. However, because paternal kin share paternal, not maternal genes, this maternal behavior cannot be explained as a standard form of kin selection, which requires relatedness between an actor (here the mother) and a non-descendant recipient (here the similar-looking infant). To address the question of how and whether such a behavior may evolve, we formally demonstrate an evolutionary mechanism by which mothers may gain fitness benefits from favoring nepotism between their own offspring and their offsprings’ PHS, as infants themselves benefit from interacting repeatedly with kin. The mechanism that we propose to call “second-order kin selection”, can be generalized beyond mother-infant interactions. While it is a novel explanation for the evolution of nepotism, second-order kin selection perfectly fits to the mathematical framework offered by inclusive fitness theory^1,24^.

## Results and Discussion

### Empirical evidence in mandrills

#### 1. Increased resemblance among PHS infants

We first show that PHS infants resemble each other more than MHS and NK do. We retrained VGG-Face, a popular algorithm for human face recognition, to recognize 112 individual mandrills from their face, independently of their position, lighting or facial expression, using approximately 12k training pictures. These pictures were taken in the course of a long-term field project on a large social group of mandrills in Gabon. We used this retrained model to predict facial distance (calculated within the encoding space of the model, see Materials and Methods) among 80 individually-recognized infants (0-1 yr) of both sexes (39 females, 41 males), represented by a total of 204 new pictures (1-10 pictures/infant), not used for training the model. Maternal identity was known for all 80 infants based on direct observations. Due to field constraints, paternal identity was determined genetically for 15 infants. For the remaining 65 infants with unknown paternity, we differentiated pairs of infants conceived during the tenure of the same alpha male from those conceived under tenures of different alpha males. Indeed, reproductive skew in mandrills is high, with alpha males monopolizing 60-70% of all conceptions^17,18^. While infants born during the tenures of “different alpha males” are thus most likely not paternally related, those born during the tenure of the “same alpha male” presumably include a large proportion of PHS. Using a linear mixed model (LMM) on this large dataset (N=2,616 dyads), and controlling for infant age and sex as well as maternal identity, we found that infants born during the tenure of the same alpha male or that were sired by the same genetic father resemble each other significantly more (lower average facial distance; Figure 1, Table 1) than those born during the tenures of two different alpha males or to two different fathers (higher average facial distance). In contrast, at these young ages, MHS do not look more alike than infants born to different mothers (Table 1). Importantly, infants born during the tenure of the “same alpha male” include dyads of infants sired by different males (~30-40%), we therefore expect PHS infants to resemble each other even more than what we report here. This result therefore strongly supports and extends one of our previous key results in mandrills: starting as early as infancy (this study) and continuing throughout juvenescence and adulthood for females^17^, PHS resemble each other more than MHS do.

**Fig. 1.**
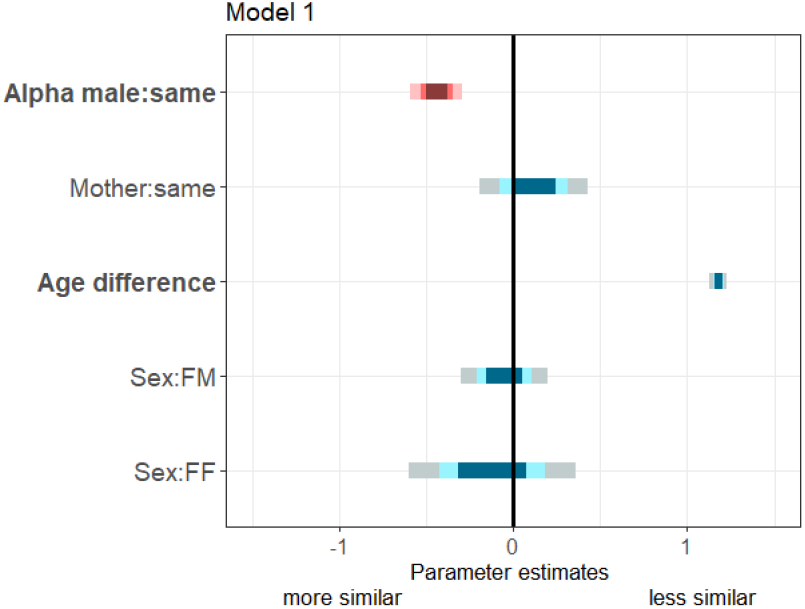
Summary of fixed effects parameters included in the model analyzing the facial distance among infants. For each estimate, the 50% (inner), 70% (middle) and 95% (outer) Wald confidence intervals are shown. Pink shades highlight the variable of interest while blue shades correspond to control variables. The following variables are displayed: infants born from different vs. same alpha male tenures or fathers (“Alpha male:same”; reference category: different); infants born from different vs. same mothers (“Mother:same”; reference category: different); infants’ difference in age (“Age difference”); infants’ difference in sexes (“Sex:FM” and “Sex:FF”; reference category: MM). Bold y-axis labels highlight variables with significant effects (P<0.05).

**Table 1.** Predictors for facial distance among infants. Significant predictors (P<0.05) are in bold (LMM with a Gaussian response).

|  |  | Infants' facial distance<br>N = 2,616 dyads |  |  |  |
| --- | --- | --- | --- | --- | --- |
| | | $\chi^2$ | $P > \chi^2$ | Estimate | Standard error |
| Predictors | **Alpha male/father at infants' conception<sup>a</sup>** | 25.70 | **<0.001** | -0.44 | 0.09 |
|  | Infants' mother(s)<sup>b</sup> | 0.39 | 0.53 | 0.12 | 0.19 |
|  | **Infants' difference in age** | 1272.2 | **<0.001** | 1.18 | 0.03 |
|  | Infants' sex<sup>c</sup> | 0.19 | 0.91 | ff: -0.12;<br>fm: -0.05 | ff: 0.29;<br>fm: 0.15 |

<sup>a</sup>Reference: different alpha males (or different fathers)
<sup>b</sup>Reference: different mothers
<sup>c</sup>Reference: mm (male-male)

#### 2. The driving role of mothers

Next, we show that mothers guide their offspring’s social opportunities towards similar-looking infants and that this effect is robust while several other predictors of offspring associations are simultaneously controlled for.

We restricted our image data set to those infant dyads for which we had detailed records on spatial associations routinely collected during behavioral observations (N=48 infants and their 30 mothers). Using generalized linear mixed models (GLMM with negative binomial family for over-dispersed count data), we analyzed spatial associations i) among infants (“infant-infant”, N = 282 dyads); ii) between infants and other mothers (“mother-infant”, N = 580); and iii) among mothers having infants at the same time (“mother-mother”, N = 325) as a function of the residuals (see Materials and Methods) of facial distance among infants, and controlling for matrilineal origin, maternal relatedness and rank and infant age and sex.

We found that mothers associate significantly more with infants that look like their own infants compared to infants that do not (Figure 2, Table 2). Infants also associate significantly more with other infants that look like them more. In contrast, associations among mothers are not influenced by the average facial distance of their offspring (Figure 2, Table 2), highlighting the fact that mothers actively associate with similar-looking infants but not with their mothers, although the latter are probably also spatially close given mother-offspring association patterns (Supplementary Information S4). This result further indicates that possible pre-existing friendships among mothers, or among these mothers and a common father (Supplementary Information S2a), are not responsible for the relationships observed between mother-infant and infant-infant dyads.

**Fig. 2.**
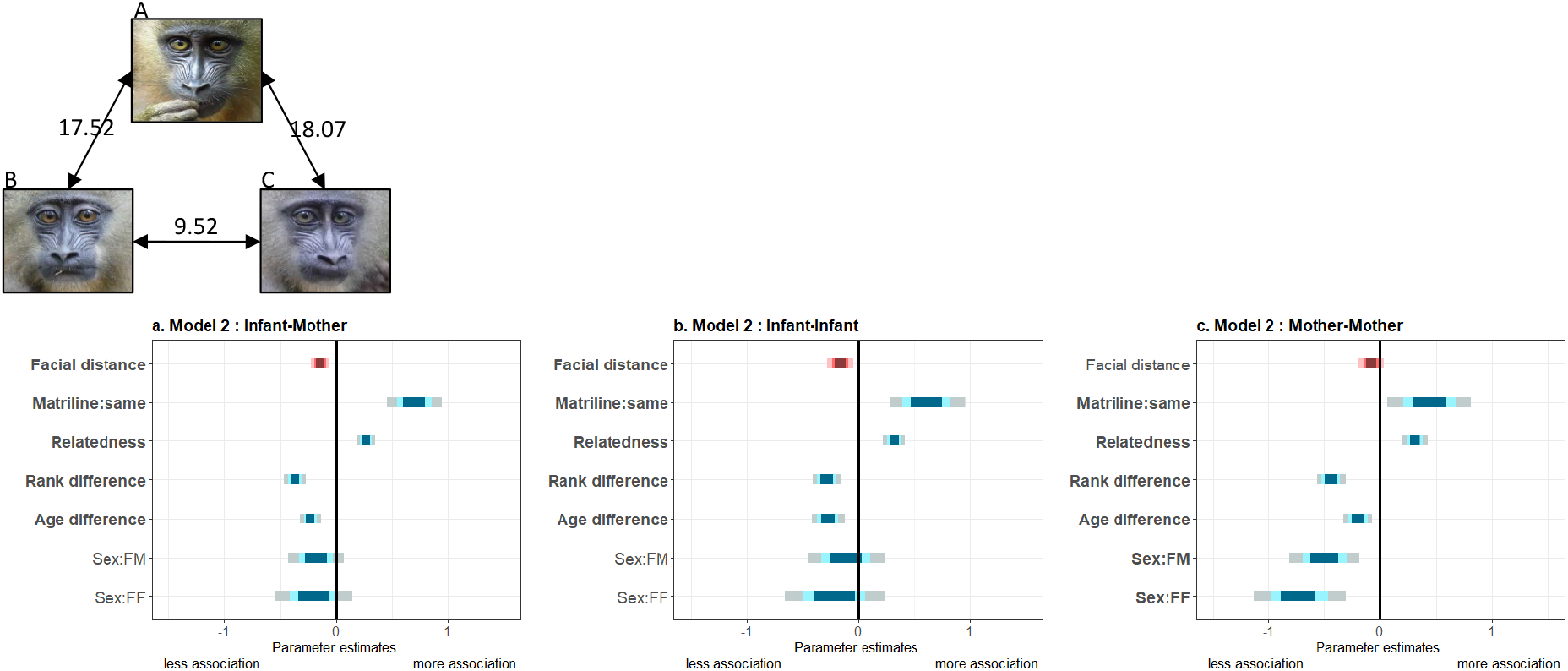
Summary of fixed effects parameters included in the three models analyzing spatial association across dyads of a. mothers and other infants; b. infants; and c. mothers. For each estimate, the 50% (inner), 70% (middle) and 95% (outer) Wald confidence intervals are shown. Pink shades highlight the variable of interest while blue shades correspond to control variables. The following variables are displayed: infants’ residuals of facial distance (“Facial distance”); different vs. same mothers’ matriline (“Matriline:same”; reference category: different); mothers’ relatedness (“Relatedness”); mothers’ difference in rank (“Rank difference”); infants’ difference in age (“Age difference”); infants’ difference in sexes (“Sex:FM” and “Sex:FF”; reference category: MM). Bold y-axis labels indicate variables with significant effects (P<0.05). Pictures depict three male infants with their average facial distances: B and C resemble each other most, in contrast to A.

**Table 2.**
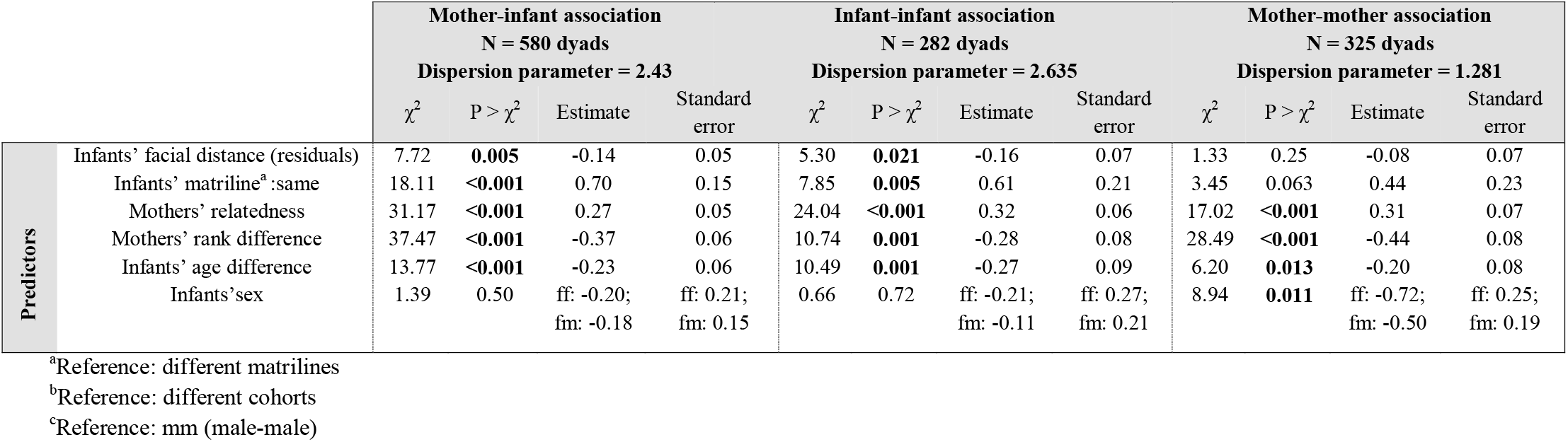
Predictors of the spatial associations recorded across mother-infant, infant-infant and mother-mother pairs. Significant predictors (P<0.05) are in bold (GLMM with negative binomial response family and log link). The reported dispersion parameter is the so-called shape parameter of the negative binomial distribution.

In mandrills, mothers constitute the main social partner for their offspring for several months or years. During their first year of life, infants are closely associated with their mother (Supplementary Information S4). To confirm that mothers, and not infants, drive patterns of association observed, we analyzed the directionality of mother-offspring follows and approaches. We found that infants follow and approach their mothers in 97.6% and 89.4% of these events, respectively (follow: N=2,387 total events recorded on 139 mother-infant dyads; approach: N=4,482 total events recorded on 155 mother-infant dyads). Therefore, in the vast majority of cases, infants go, indeed, where their mothers are or go.

Although we do not assume this process to be conscious, mandrill mothers associate significantly more often with infants that look more like their own offspring, possibly orientating their offspring’s social preferences towards similar-looking other infants that are also more likely to be PHS than a random pair of infants. From rodents to humans, mothers often show some form of control over social opportunities and preferences of their offspring, and mammalian mothering may impact both their neural and social development^25^. In chimpanzees, mothers with sons are more gregarious and associate more with adult males than mothers with daughters, thereby determining their sons’ social trajectories^26^. Despite strong maternal control over offspring’s social choices in mammals^25,27^, we explore below an alternative scenario where fathers, not mothers, may use prominent visual resemblance among PHS to target paternal care and mediate PHS nepotism, thereby ultimately favoring the transmission of paternal genes.

#### 3. Alternative hypothesis: Is increased PHS resemblance driven by father-based selection?

To care for their own offspring, mandrill fathers need some form of paternity certainty because females mate with several males around ovulation^28^. In some other promiscuous species, true paternal care has been unambiguously demonstrated: male baboons, for example, selectively support their own offspring in agonistic disputes^29^. If mandrill fathers know with certainty at least one or a few of their offspring, then increased facial resemblance may help discriminating others. This mechanism, although plausible, is subject to cognitive constraints and a precise knowledge about female fertility as female mandrills are only fertile during a restricted time window (May-Sept^30^). More parsimoniously, mothers may remember whom they mated with and associate with the putative father after birth to promote paternal care. Heightened facial resemblance among PHS could then help fathers to discriminate all of their offspring from other infants. Contrary to male baboons, however, paternal care in mandrills is elusive: males neither carry infants nor do they groom or affiliate with them (MJEC pers. obs.), even though in captive colonies, males are spatially closer to their own juvenile offspring than to unrelated juveniles of similar age^19^. While we cannot exclude paternal care as a driving force for increased facial resemblance among PHS, patterns of male residency in this natural mandrill population do not provide strong support for this scenario. Indeed, males are mainly present during the short breeding season, although some males remain in the group for varying periods of time, from a few days to a few months^31^. Among 69 male.years (N=29 different subordinate and dominant males) observed throughout the study period (8 reproductive seasons), only 44.7% and 54.5% of males, respectively, that were present during a given reproductive season remained in the study group until the next birth season. Opportunities for fathers to personally care for their infants therefore occur statistically only half of the time. Finally, those males that were present during a given reproductive cycle are not responsible for maintaining proximity with either infants or their mothers (MJEC and BRT, pers. obs.).

### The mother’s benefits of matching her infants with related ones

Because PHS share paternal, not maternal genes, we finally show how, under minimal assumptions, mothers can obtain fitness benefits by fostering interactions between their offspring and paternally-related siblings. Key steps to that conclusion are that infants engage in repeated interactions within dyads, and that such interactions are mostly of two types, “repeatedly cooperate” or “repeatedly defect”, so that only dyads of cooperators can repeatedly cooperate. Then, any maternal behavior that increases the likelihood that her offspring will repeatedly cooperate rather than repeatedly defect may be favored. This two-step reasoning does not require a statistical association (linkage disequilibrium) between alleles affecting the control of assortment of infants by mothers, and alleles affecting cooperation. To reach this conclusions, we will use a “one-generation” formalism^32^ suitable to take into account both the interactions between relatives and multilocus processes, and which has proven useful in particular to avoid double counting of fitness benefits (as may happen when compounding fitness effects of related individuals over different generations). In the present case, it allows to correctly account for fitness benefits to mothers when infants are involved in social interactions.

In this formalism, selection on an allele affecting the mother’s fitness is quantified as a covariance ^33^ between an indicator variable for presence of the allele in a mother, and mother’s fitness. This fitness is her number of (adult) offspring, expressed as a function of her own behavior and of that of her social partners (neighbor-modulated fitness), and then as a function of gene effects underlying such behaviors. The covariance expression for selection then involves expected values of products of the mother’ s indicator variable with the variables describing gene effects in different individuals, that can be interpreted in terms of relatedness coefficients and linkage disequilibria. In the present case, we will see that many such products can be ignored, allowing a simple characterization of selection on maternal behavior.

For simplicity, we assume effects on infant survival. We could alternatively assume effects on infant reproductive potential (“quality”) but then the fitness of a mother should be measured as a weighted sum of numbers of offspring of different quality, which would complicate the model formulation without modifying its conclusions. Likewise, formal models including pairing processes are generally complex to formulate but such a formulation is not required to understand the key qualitative features of the present scenario.

First, consider the fitness benefits for paired infants. Let us assume that pairs of infants play an iterated prisoner’s dilemma in its canonical form (with *R* indicating the “rewards” received by two cooperating infants, *P* the “punishment” payoff received by two defecting infants, *T* the “temptation” payoff received by a defecting infant interacting with a cooperating infant, and *S* the “sucker’s” payoff received by a cooperating infant interacting with a defecting infant, and *w* the probability of iteration^34^, and a tit-for-tat strategy in response to defection. Accordingly, after multiple interactions the expected payoffs are *R*/(1 – *w*) for pairs of cooperators, *P*/(1 – *w*) for pairs of defectors, and respectively *S* + *Pw*/(1 – *w*) and *T* + *Pw*/(1 – *w*) for a cooperator and a defector paired together. Provisionally assuming that cooperators and defectors are equally frequent, the average payoff is therefore (*R* + *P*)/[2(1 – *w*)] for identical dyads, and ((1 – *w*)(*S* + *T*) + 2*Pw*)/[2(1 – *w*)] for non-identical dyads. For long-lasting interactions (*w* → 1), the relative values of the payoffs of identical and non-identical dyads compare essentially as *R* + *P* vs. 2*P*. Given that cooperation is mutually beneficial (*R* > *P*), identical dyads are favored over non-identical ones. If cooperators and defectors are not equally frequent, the average payoff of identical dyads will scale as a weighted average of *R* and *P* but the reasoning and main conclusion is otherwise unchanged: identical dyads still enjoy an average fitness benefit proportional to *R* – *P*. This conclusion embodies the fact that, on average over cases where they cooperate and cases where they do not, individuals can benefit from increasing the likelihood of interacting with identically-behaved ones, whether relatives of not. Therefore, they can benefit by increasing the likelihood of interactions between relatives, and it would readily explain kin recognition by infants, which we do not assume here.

Next, consider selective effects on alleles acting in mothers to control the assortment of their infants. For simplicity, let us assume that payoffs affect infant survival, so that the mother’s fitness is proportional to the survival probability of her offspring. We can write the expected payoff (and, up to a constant factor, the linearized survival benefits) for a focal infant as *W* ≔ *β*_0_ + *β_z_z* + *β_z_p__z_p_* + *β_zz_p__zz_p_*, where *z* is the indicator variable for the event that the focal infant initiates the interaction by cooperating, *z_p_* is the same variable for its partner, and the *β*s are functions of the model parameters. Then, if a “mutant” allele increases by *δ*, relative to a “resident” allele, the likelihood that infants assort in identical pairs, this mutant will experience a total fitness effect proportional to *δΔW*, where *ΔW*: = *β_z_Δ*E(*z*) + *β_z_p__Δ*E(*z_p_*) + *β_zz_p__Δ*E(*zz_p_*) where for any variable *x, Δ*E(*x*) is the difference in expected value of *x* between infants of mothers bearing the mutant vs. those of mothers bearing the resident allele.

We first focus on the last term of the selective effect *ΔW, β_zz_p__Δ*E(*zz_p_*), as we will later see that other terms are comparatively negligible. This represents the fitness effect of the interaction of events represented by *z* and *z_p_*. The system of four equations, implied by the expression for *W* for all combinations of the two indicator variables, shows that *β_zz_p__* is the difference between the unweighted average payoff of identical dyads (z and z_p_ equal to each other) and the unweighted average payoff of non-identical dyads. As shown above, this difference is proportional to *R* – *P* > 0. Consider further that phenotypes are affected by the additive effect of the alleles transmitted by the parents, *z* = *C_m_* + *C_f_* where *C_m_* and *C_f_* are effects inherited from mother and father, respectively. Likewise, for the infant partner, *z_p_* = *C*_*m*(*p*)_ + *C*_*f*(*p*)_ in terms of effects *C_m_*(*_p_*) and *C*_*f*(*p*)_ inherited from the partner’s mother and father. Then the survival of an offspring will increase with any of the products *C_m_C*_*m*(*p*)_, *C_m_C*_*f*(*p*)_, *C_f_C*_*m*(*p*)_ and *C_f_C*_*f*(*p*)_.

We do not assume any relatedness among mothers (which would increase the expected value of *C_m_C*_*m*(*p*)_), nor do we assume any form of (dis-)assortative mating increasing or reducing the expected value of cross-sex products. Then, only *δβ_zz_p__Δ*E(*C_f_C*_*f*(*p*)_) remains, meaning that any maternal behavior that increases the expected value of *C_f_C*_*f*(*p*)_ over offspring would enjoy fitness benefits. The conditions under which such benefits may outweigh costs possibly associated with the choice process are general conditions favoring choice: a high variance in quality (here, a high variance of inherited effects on cooperative behavior), and a strong impact of choice on expectation of *C_f_C*_*f*(*p*)_. which is dependent on a high variance in male reproductive success within a cohort of infants (the case in mandrills^18,35^) and on the existence of cues to infer paternal relatedness between infants, here increased facial resemblance among PHS.

This fitness effect results from the fact that infants being more similar at loci controlling phenotypic similarity are more similar at loci controlling cooperation. Such shared similarity (“identity disequilibrium”) automatically results from an increased likelihood of shared paternity. It is expected, indeed, in any population where there is variation in relatedness among individuals, even in a finite panmictic population^36^. When some pairs of individuals are more related to each other than other pairs, the fact that a pair is identical at one locus is an indication that the pair is more related than a random pair on average, and then, it also tends to be more identical than random pairs at other loci in the genome^37^. By contrast, *δΔ*E(*z*) may be positive only if the alleles affecting the control of assortment of infants become statistically associated to cooperator alleles in mothers’ genotypes. Such linkage disequilibria are typically of lower magnitude than identity disequilibrium, as recombination is generally efficient in reducing them^38^. Likewise, *δΔ*E(*z_p_*) depends on a statistical association between the alleles affecting the control of assortment and cooperator alleles borne by the infant partner, and given that we do not assume that mothers are able to recognize any cooperator allele in any infant, such an association is all the more likely to be weak.

Although this mechanism does not assume any increased relatedness among mothers of interacting dyads over mothers of non-interacting dyads, such variation in relatedness may be difficult to fully exclude in natural populations. If it is present, a selection effect *δβ_zz_p__Δ*E(*C_m_C*_*m*(*p*)_), analogous to the above one, arises and represents a standard kin selection effect since actor and recipient are then related. Yet this kin-selection effect arises only in conditions where the previous effect arises, and will be comparatively small when maternal relatedness is small relative to the relatedness between paternally-derived gene copies. Thus, even if present, a kin-selection effect will be small compared to the main selection effect favoring assortment between paternal half-sibs.

The main conclusion is therefore that selection on mother’s control of assortment of infants is proportional to *R* – *P* and to the identity disequilibrium she creates by such control. By favoring nepotism between their own offspring and their PHS, mothers would therefore derive direct fitness benefits (Figure 3, scenario A). The mechanism described here can be generalized to any actor whose behaviors promote positive interactions between a non-kin recipient and any actor’s kin (Figure 3, scenario B). We term the mechanisms described in this generalized version “second-order kin selection”.

**Fig. 3.**
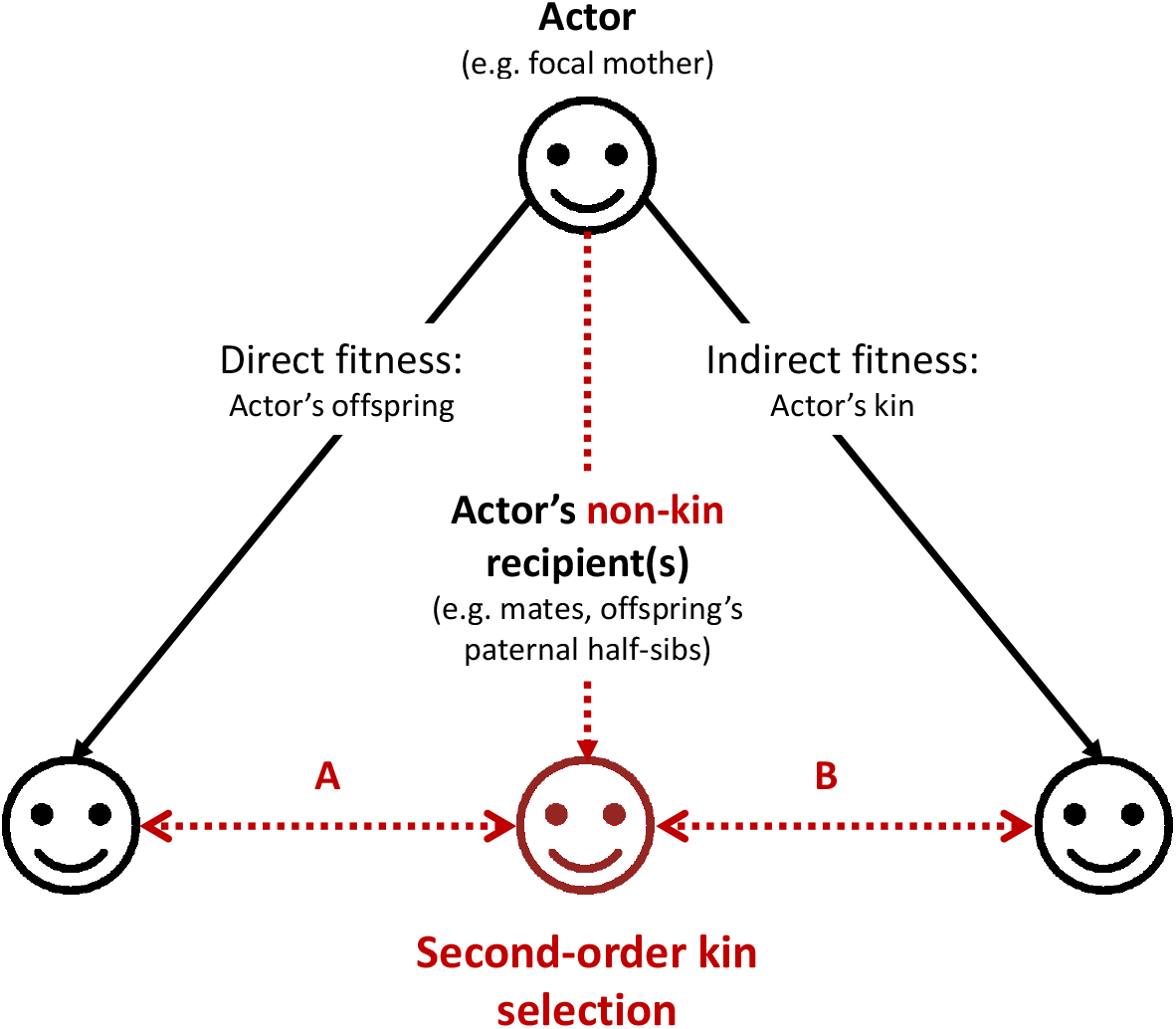
Graphical representation of the “second-order kin selection” process. The actor (here a focal mother, face on the top) has the control over her behaviors with social partners (all three faces on the bottom including actor’s offspring, other kin and non-kin in red). Plain black arrows represent the inclusive fitness framework of the kin selection theory. Dashed red arrows represent the second-order kin selection process where an actor’s social behaviors towards a non-kin recipient (e.g. an offspring’s paternal half-sib) favor the latter’s social behaviors towards the actor’s kin (offspring or other kin). We provide a few examples (A: the mechanism explored in this study; B: a generalization of the mechanism to other actor’s kin) where this process may occur (and see discussion). Importantly, the second-order kin selection necessitates that non-kin recipient in red shares genetic or reproductive interests with the actor’s kin (double-arrows).

## Conclusion and Future Directions

Given mandrills’ biology, the most plausible interpretation of the heightened facial resemblance observed among paternally related individuals, from infancy to adulthood^17^, is based on the notion that this resemblance has been selected to allow mothers to foster interactions between their offspring and their offspring’s paternal relatives. We propose the term “second-order kin selection” to explain such behavior. This process is, indeed, distinct from classic kin selection as actors (mothers) may derive direct fitness benefits from interacting with non-kin recipients (their offspring’s paternal half-sibs) because of the benefits of these interactions towards the actors’ own kin that occur through the increased expected payoffs of reciprocal interactions between kin.

Two critical conditions of this process are the ability to predict relatedness between assorted individuals (here by phenotype matching) and the occurrence of reciprocal interactions between the matched social partners. We have provided evidence that the first condition is met in mandrills. In particular, we have both shown that relatedness is signaled and that these signals modify the behavior of receivers (mothers). A firm demonstration of the second condition will necessitate years of detailed behavioral and demographic data and is beyond the scope of this study. However, reciprocal interactions have been well demonstrated in other study systems. Females baboons, for example, form the most enduring and reciprocal social bonds with their close kin^39^. In addition, the positive and strong relationship between individual fitness and the quality of the social environment, often approximated using measures of spatial association, is no longer in question. Indeed, social integration is one of the strongest predictors of health and longevity in humans^40,41^. This conspicuous relationship is also observed in many social mammals – with effect sizes of strikingly high amplitudes^42^. In all mammalian orders, individuals that are poorly social integrated display considerably shorter life spans and decreased reproductive success than individuals enjoying a rich social life^43–47^.

To the best of our knowledge, this form of selection has been overlooked so far and how this process shapes the social structure of animal societies is still unknown. Yet, cooperation among non-genetically related in-laws (affinal kin), often reported in humans^48–50^, fits into the second-order kin selection process we propose because affinal kin or spouses may share genetic interests in the next generation (see also^51,52^). In nonhuman animals, the generality of this process will depend on the taxonomic distribution of reciprocal interactions. Other modulating factors involve the variance in male parenting, which influences the benefits for a kin recognition mechanism, the conditions that the matched individuals are unable to perform this recognition themselves, and the opportunity for control of assortment of relatives. Several of these conditions may be widespread in animal societies. Although the opportunity for control of kin assortment may be the most restrictive condition, mothers guide their offspring’s social development in many social mammals^23,25^ and other conditions similarly leading to direct selection for assortment of related offspring could operate elsewhere, such as in the various forms of cooperative breeding or nest parasitism in birds. Finally, our study highlights how state-of-the art artificial intelligence algorithms combined with long-term field data can help unravelling adaptive details of social complexity that would otherwise go completely unnoticed.

## Materials and Methods

### Study population

Since 2012, we have been monitoring the only habituated, free-ranging and un-provisioned group of mandrills, living in Southern Gabon (Lékédi Park, Bakoumba) within the framework of the “Mandrillus Project”^17,31^. This group originated from 65 individuals that initially lived in a semi-free ranging population housed at CIRMF (Centre International de Recherches Médicales de Franceville, Gabon) and released on two occasions into the park (2002 and 2006^53^). Captive-born females reproduced with wild immigrant males starting the first year post-release. In 2020, the group was composed of ca. 250 individuals of both sexes and all ages, about 200 of them being individually known and monitored on a daily basis; wild-born individuals represented about 95% of the animals studied for this project.

During daily observations, we record detailed data on group composition and behavior. In addition, we take pictures of unambiguously identified individuals when visible and close (less than 10m away). In this study, we considered a total of 80 infants (39 females, 41 males; aged 4-365 days) who contributed to the dataset of portrait images and 48 infants (24 females, 24 males) born from 30 different mothers contributing to the study of spatial associations as a function of facial resemblance (excluding those infants that were aged more than a year apart or those for whom no behavioral data was available). Dates of birth were exactly known for 31 infants and estimated within a time-window of one day to two months for the remaining 49 infants based on patterns of mother’s sexual swellings and infants’ physical appearance.

## Measuring facial resemblance

### Image database and pre-processing

The mandrill face database includes ~16k images representing 276 different mandrills originating from the study natural population (12.9k images), a semi-captive population housed at CIRMF (Centre International de Recherches Médicales de Franceville, Gabon; 2.7k images) and other sources (the Internet, the Wildlife Reserves of Singapore, Zoo of Grandy; 0.4k images). Images from Gabon (natural and CIRMF populations) were taken between 2012 and 2018. Pictures represent individuals that are awake and passive, awake and active (i.e. feeding, grooming, vocalizing) or anesthetized during several captures that occurred between 2012 and 2015 (representing 1.1k images). We frequently photographed active individuals using the slow burst mode of cameras, which allows to capture variation in face position and expression while avoiding identical frames. The multiple frames obtained when using the slow burst mode are hereafter referred to as a “burst-mode series”. Images were then manually oriented to align pupils horizontally, and cropped to generate square portraits centered on the nose and excluding the ears.

For training the DNN for both face identification and face verification, we used a learning dataset of 12.2k pictures representing all individuals of the mandrill face database >1 yr, plus infants ≤1 yr not belonging to the 80 studied infants. Based on preliminary tests described previously^17^, we included in the learning set all pictures displaying a face in frontal view (approx. <30°) and with less than 50% of occlusion, of whatever quality provided that field assistants individually recognize the animal. Because the face of a given individual varies considerably between its different age classes (infant, juvenile, adolescent -only for males-, subadult -only for males- and adult), we used “ind-age” classes rather than individuals for the identification task, that is, we treated two ind-age classes representing the same individual as distinct and independent classes. The final learning dataset contains 168 ind-age classes. This learning set was split further into a training set and a validation set. The validation set was used to parameterize the model for the face identification task. It contained two images of each class. For a correct evaluation of training performances, we ensured that none of the validation image was from a burst-mode series that also contained images present in the training set. The training set contained all other images of the learning set. Because we were able to reach high performances despite a large imbalance between classes in the training sets, we did not attempt to correct for this imbalance.

The test dataset includes images of infants (≤1 yr) from the study natural population. To maximize the quality of dissimilarity measurements, we selected only the best-quality pictures: face in frontal view, sharp image, neutral light, occlusion limited to below the mouth, neutral facial expression, no shadow or lighting spot, single image of a burst-mode series. Eventually, we analyzed 204 images representing 80 different infants (mean ± SD = 2.55 ± 1.79 pictures per infant). For the sake of clarity, we alternatively use expressions like “facial resemblance” (facial similarities) in the main text or “facial distance” (facial dissimilarities). In other words, pairs of infants that look like each other present, on average, low facial distances.

### Face identification

We trained a DNN to identify ind-age classes as a goal to learn a deep representation of mandrill faces. We used the popular VGG-Face^54^ as a starting point and retrained this network with mandrill portraits. This procedure, called “transfer learning”, allows to reach high model performance even with relatively small datasets^55^. VGG-face is a VGG16 that previously learned to recognize 2.6k different humans from a total of 2.6M portrait pictures. We replaced the last softmax classification layer of VGG-Face by a new layer of 168 dimensions (which corresponds to the number of classes in the new mandrill identification task). We included two dropout layers (with 50% dropout probability), one after each fully connected layer, to limit the risk of overfitting. We trained the network using a stochastic gradient descent with momentum optimizer, with initial learning rate of 10^-5^ for all convolutional layers except for the two fully connected layers and the classification layer, for which the learning rate was set to 10^-3^. The learning rate decreased by a factor 10 every 5 epochs. Learning continued until the validation loss did not decrease further after three consecutive epochs. In order to match the input size of VGG-Face, mandrill portraits were downsized to 224×224 pixels ×3 colors prior to analyses. We set the batch size to 32. We limited overfitting further by using “data augmentation”: each iteration, images were shifted horizontally and vertically (by a number of pixels randomly selected within the range [-40 40]), rotated (range of degree: [-20 20]) and scaled (range of factor: [-0.7 1.2]). The entire training procedure was repeated 10 times. Eventually, after approximately 15 epochs, VGG-Mandrill was able to identify mandrill faces individually with a maximal accuracy of 93.42% (sem: ± 0.13), and high generalization capacity (i.e. limited overfitting; Supplementary Information S5).

### Face verification

We retrained the DNN using the full learning set to maximize the number of images (i.e. with both the training and validation sets). VGG-Mandrill was then used to extract deep feature activation vectors, a compact (4,096-dimensional) and informative representation of a mandrill face. Previous experiments revealed that face verification with mandrills was most performant when using activation vectors from the first fully connected layer (after RELU nonlinear transformation), compared to using vectors from the second, or from both fully connected layers^17^. The distance between feature activation vectors predicts the resemblance between images^56^. In this study, we used a *χ*^2^ distance calculated with normalized features (see for details on the normalization step^17^).

Last, we used a linear support vector machine (SVM) to learn a distance metric, with the goal to find the feature weights that optimize face verification. We randomly selected 15k pairs of images representing different individuals and 15k pairs representing same individuals, and for each pair we calculated the *χ*^2^ difference (as in^54^). We then ran the SVM with the *χ*^2^ differences as explanatory variables and 0 (different-individual pairs) or 1 (same-individual pairs) as a response variable. The SVM outputs the accuracy of the face verification task as well as the weight of each feature. We found that the classifier could verify mandrill identity from their face with an accuracy of 86.87 % (sem: ± 0.01). Weights were eventually used to calculate a weighted *χ*^2^ distance between every pairs of images in the test set. Pairwise distances were averaged for every different pairs of individual (N = 3160 pairs of infants represented by a total of 20,421 pairs of pictures) to provide our final estimates of facial distance across pairs of infants (mean ± SD: 6.46 ± 6.99 pairs of pictures for each pair of infants).

### Behavioral observations

Trained observers perform daily behavioral observations on all individually-recognized mandrills using either *ad libitum* data or 5-min focal sampling. During each focal observation, we also record scans on one to three occasions to measure spatial associations between the focal animal and all groupmates that are located less than five meters away (termed “scan”). Since 2012, we obtained 9,305 scans from 48 infants and their 30 mothers during their offspring’s first year of life (mean ± SD: 124.1 ± 178.3 scans per individual). We restricted our analyses on association rates to infant-infant (169 ± 77 scans per pair), mother-infant (174 ± 80) and mother-mother (174 ± 91) dyads with at least 5 scans (with at least one scan for either partner) and when both individuals were alive and infants aged ≤1 yr. For each association rate, we matched the average facial distance of the corresponding pair of infants aged ≤1 yr. Here, we exclusively studied association rates and no other social behavior such as grooming because scans were more numerous than any other behavioral records, and also because we reasoned that if mothers influence their offspring’s social opportunities, this will mainly take the form of increased associations and not of an increased investment into potentially costly social behavior.

Behavioral observations further allow us to examine patterns of two conspicuous mother-infant behaviors, “follows” and “approaches” recorded as bouts during focal samplings of infants and their mothers. Finally, we reconstructed both male and female dominance hierarchies, using the outcomes of approach-avoidance interactions obtained during *ad libitum* and focal samplings and calculated using normalized David’s score (as per^17^). We divided adult females into three classes of rank of similar size across the entire study period (2012-2020; high-ranking, medium-ranking, low-ranking). For males, we used monthly rank, distinguishing the alpha male from all other subordinates. When the alpha position was highly disputed for a given month, we considered the male’s hierarchy as unclear (see also below).

#### Matrilineal identity and genetic relatedness

Maternity was known for all 80 studied infants thanks to daily monitoring. Matrilineal identity was determined for all of them, including those 48 infants and associated 30 mothers used in the proximity analyses, thanks to detailed data on births and pedigree information available since the beginning of the project^17^ and at CIRMF^18^. In particular, individuals belong to the same matriline when they share a common mother, maternal grand-mother, great-grand mother and so-on with the exception of those females born at CIRMF that belonged to the same matriline but were released on two different occasions. In these cases, major social disruptions involved changes in female’s dominance hierarchy, with females released in 2006 being automatically lower in rank than all other females released in 2002, whatever the initial rank they achieved. In these cases, females released in 2006 formed a new matriline, independent from their matrilineal origin at CIRMF.

Genetic relatedness was determined using pedigree data obtained from paternity analyses performed on most adult mandrills and on a subset of immature individuals (see for details^17^). We determined paternity for 15 of the 80 portrayed infants. For the proximity analyses, we restricted our data set to those infant dyads (and associated mother-infant and mother-mother dyads) for whom the mothers’ pedigree was known to the previous generation (maternal grand-father and grand-mother known) in order to obtain good estimate of mother-mother relatedness. Relatedness across the 285 different pairs of mothers vary from 0.01 to 0.66 (mean ± SD: 0.13 ± 0.12).

#### Statistical analyses

All data were analyzed using R v. 3.6.1. using the package spaMM.

We first fitted a General Linear Mixed Model (LMM) to study variation in the averaged facial distance among pairs of 80 infants aged ≤1 yr as a function of the averaged distance in ages across all pairs of pictures collected on each studied pair of infants, their sex (class variable with three modalities: “ff”, “fm”, “mm”), whether or not they share a common mother (class variable with two modalities: “same mother”, “different mother”), and whether or not they either share a common father or were conceived during the tenure of the same alpha male (class variable with two modalities: “same father/alpha”, “different father/alpha”). We excluded those infants (and the corresponding pairs) conceived during a month when the alpha position was disputed and unclear, resulting in 2,616 pairs of infants represented by 16,923 pairs of pictures (mean ± SD: 6.47 ± 6.74 pairs of pictures for each pair of infants). We included an autocorrelated random effect with distinct levels for each pair, to represent correlations between pairs of facial distance values involving a same infant in both distances, with a single correlation value for all such pairs. Its correlation is equal to 1 between permuted pairs (which are thus affected by the same value of the random effect), equal to a correlation value ρ (fitted to the data) between pairs sharing one individual, and to 0 for non-overlapping pairs. The case where ρ equals 0.5 represents the case of two additive individual random effects, each with the same variance. The fitted correlation model is therefore more general than this particular case. It is formally identical to a random effect model for a “diallel” experiment as considered in quantitative genetics, where the same individual may be the “first” or the “second” member of a mating pair across different mating pairs, and more specifically assuming that in each case it would express the same effect on its offspring (implying that there is a single variance parameter to be fitted). The present random effect structure accounts for the analogous symmetry of effects on facial distance. The case where ρ=0.5 represents additive random effects from each individual (analogous to “General combining abilities” in quantitative genetics^57^) and deviations from ρ=0.5 additionally represent non-additive effects (“specific combining abilities”). We fitted a heteroscedastic residual variance with prior weights defined as the total number of pairs of pictures collected on each dyad of infants (giving more weight to those pairs with more numerous pictures). We standardized the averaged distance in ages to allow comparisons with other estimates.

We then fitted three Generalized Linear Mixed Models (GLMM) with negative binomial response family and log link to assess predictors of the frequency of spatial association among dyads of i) infants (“infant-infant”); ii) infants and other mothers (“mother-infant”); and iii) mothers having infants in the social group (“mother-mother”). The response is the number of observed associations. Each fitted model includes an offset (the logarithm of the total number of scans recorded on both individuals of the focal pairs), six fixed effects, a random effect term representing the effect of two additive individual random effects under the constraints of symmetry, as for “general combining ability” in a “diallel” experiment, in models of symmetrical interactions (mother-mother and infant-infant), and standard individual-level random effects for mothers and for infants in the mother-infant model. The six fixed effects are whether the infants belonged to the same matriline (class variable with two modalities: “same matriline”, “different matriline”), relatedness between mothers, mothers’ rank difference (continuous variable with three values: “0” for no rank difference, “1” for one rank difference, for example, between high-ranking and medium-ranking mothers; “2” for two rank differences, for example, between high-ranking and low-ranking mothers), infants’ age difference when behaviors (not photographs) were recorded (continuous variable in days and <365), infants’ sex (class variable with three modalities: “ff”, “fm”, “mm”), and the residuals of infant facial distance. Indeed, to assess the effect of infant facial distance unconfounded by the effect of infant age differences (at the time of pictures’ collection), we used the residuals of such distance obtained from the fit of an LMM as performed above and that included as predictor the age difference (averaged distance in ages across all pairs of pictures), and an autocorrelated random effect with the same “diallel” correlation structure as above (N = 3160 pairs of infants, represented by 20,421 pairs of pictures).

Both matrilineal origin and females’ relatedness were independent from facial distance values meaning that highly resembling infants were not more likely to be found within the same matriline than lowly resembling infants nor were they the offspring of highly related females (e.g., “infant-infant” data set: Pearson correlation between facial distance and mothers’ relatedness: r = 0.008, P=0.89; facial distance within same *vs*. different matriline, mean ± SD: 13.03 ± 2.12, N = 56 pairs of infants *vs*. 13.21 ± 1.90, N = 226 pairs of infants; and see Supplementary Information S6).

We standardized all continuous predictors to allow comparisons of the estimates and we preliminary checked that no significant first-order interaction occurred among all these explanatory variables (not shown) and kept full models as final models. In all these three GLMM, we further checked for possible hazardous multicollinearities between continuous predictors by calculating variances of inflation (VIF < 2). For the two significant negative effects of facial-distance residuals, we excluded individuals one at a time and checked that there was no apparent error in the data for individuals most supporting the negative relationships. To validate the models, we finally verified that the magnitude of standardized residuals is independent of the fitted values.

## Acknowledgments

We are grateful to past and present field assistants of the Mandrillus Project who collect daily behavioral data on the study population, to the Wildlife Reserves of Singapore and the Zoo of Granby and to the Primatological Centre at CIRMF (Gabon) for providing pictures of their mandrills, to the SODEPAL-COMILOG society (ERAMET group) for their long-term logistical support and contribution to the Mandrillus Project. This study was funded by several grants that allowed long-term collection of data: Deutsche Forschungsgemeinschaft (DFG, KA 1082-20-1) to PMK and MJEC, SEEG Lékédi (INEE-CNRS) to MJEC, and Agence Nationale de la Recherche to MJEC (ANR SLEEP 17-CE02-0002) and to JPR (ANR-20-CE02-0005-01). This study was approved by an authorization from the CENAREST institute (permit number: AR017/22/MESRSTTCA/CENAREST/CG/CST/CSAR). This is a Project Mandrillus publication number ## and ISEM ##.

